# Variable oxygen environments and DNMT2 determine the DNA cytosine epigenetic landscape of *Plasmodium falciparum*

**DOI:** 10.1101/2021.12.09.471905

**Authors:** Elie Hammam, Samia Miled, Frédéric Bonhomme, Benoit Arcangioli, Paola B. Arimondo, Artur Scherf

## Abstract

DNA cytosine methylation and its oxidized products are important epigenetic modifications in mammalian cells. Although 5-methylcytosine (5mC) was detected in the human malaria parasite *Plasmodium falciparum*, the presence of oxidized 5mC forms remain to be characterized. Here we establish a protocol to optimize nuclease-based DNA digestion for the extremely AT-rich genome of *P. falciparum* (>80% A+T) for quantitative LC-MS/MS analysis of 5-hydroxymethylcytosine (5hmC), 5-formylcytosine (5fC) and 5-carboxylcytosine (5caC). We demonstrate the presence of 5hmC, 5fC and 5caC cytosine modifications in a DNMT2-only organism and observe striking ratio changes between 5mC and 5hmC during the 48-hour blood stage parasite development. Parasite-infected red blood cells cultured in different physiological oxygen concentrations revealed a shift in the cytosine modifications distribution towards the oxidized 5hmC and 5caC forms. In the absence of the canonical C5-DNA methyltransferase (DNMT1 and DNMT3A/B) in *P. falciparum*, we show that all cytosine modifications depend on the presence of DNMT2. We conclude that DNMT2 and oxygen levels are critical determinants that shape the dynamic cytosine epigenetic landscape in this human pathogen.

## Introduction

Since its discovery in 1948^1^, 5-methylcytosine (5mC) in DNA, also known as the 5th base of the genome, is now the best characterized epigenetic mark in higher eukaryotes, playing very important functions in different cellular processes such as cell differentiation and X chromosome inactivation^2,3^. DNA cytosine methylation is catalyzed by members of the DNA methyltransferases family (DNMT1 and DNMT3A/B)^4–6^ and 5mC is mostly found in CpG rich regions of gene promoters (CpG islands or CGIs)^7–9^. The presence of 5mC within CGIs of promoter regions is often linked with the inactivation of the corresponding gene^10,11^. This occurs through the impairment of the binding of transcription factors and the recruitment of methyl CpG binding proteins (MBDs) creating a transcriptionally repressive environment^12^.

Besides 5mC, another major DNA cytosine modification in eukaryotes is 5-hydroxymethylctosine (5hmC)^13^. 5hmC is generated as the first oxidative product of the TET (Ten eleven translocation)-dependent DNA cytosine demethylation pathway^14^. TET enzymes (namely TET1, TET2 and TET3 in mammalian cells^15^) progressively oxidize 5mC into 5hmC and further into 5fC (5-formylcyotsine) and 5caC (5-carboxylcytosine)^15^. These latter can be actively removed by different mechanisms including the activity of a thymine DNA glycosylase (TDG) coupled with the base excision repair machinery (BER) or passively lost upon DNA replication^14^. It is now well established that 5hmC is on its own an independent epigenetic mark in particular in the brain^16,17^. In addition, it has been suggested that 5fC and 5caC may also have regulatory functions especially when present at the promoters of highly expressed genes^18,19^. Importantly, TET proteins require cofactors for their activity such as alpha-ketoglutarate, iron and oxygen (O_2_)^20^ The latter is a very important factor that modulates the activity of TET enzymes such as in tumours^21,22^.

DNA cytosine modifications have also been reported for the extremely AT-rich genome (>80%) of the protozoan malaria parasite *P. falciparum*^23,24^. This important human pathogen is exposed to fluctuating oxygen levels during its complex life cycle in the human host (liver and blood) and *Anopheles* mosquito vector (midgut and salivary glands). While circulating parasite infected erythrocytes (IE) are exposed to different O_2_ levels in patients that vary between 5% in venous blood and 13% in arterial blood^25^, these levels can reach up to 20-21% in the mosquito vector, where the tracheal system enables transport of gaseous oxygen from the atmosphere directly to the inner organs^26^. Fluctuating O_2_ levels could constitute a sensor system which participates in a rapid adaptation of *P. falciparum* to its immediate cellular environment.

In this study we analyzed if different oxygen level environments encountered during the life cycle of *P. falciparum* can impact DNA cytosine methylation, 5mC, and its oxidative derivatives, 5hmC, 5fC and 5caC. To this end, we used ultra-performance liquid chromatography-tandem mass spectrometry (LC/MS-MS) on digested genomic DNA for the identification and quantification of 5mC and its oxidative derivatives for an extremely AT-rich genome. We showed that DNA cytosine 5mC/5hmC ratio is dynamically regulated throughout the 48-hour asexual blood stage parasite progression with a sharp increase of 5mC at the trophozoite stage. Importantly, hyperoxic culture conditions led to a dramatic shift of DNA cytosine modifications towards the oxidative derivatives 5hmC and 5caC. Given the absence of the DNA methyltransferase family genes DNMT1 and DNMT3A/B, malaria parasites belong to a small group of DNMT2-only organisms^27^. *P. falciparum* DNMT2-mediated tRNA methylation is not essential during standard *in vitro* culture conditions but was shown to maintain cellular homeostasis in the presence of different stressors^28^. In this work we establish that plasmodial DNA cytosine modifications depend on the presence of DNMT2. The observed dual specificities of this enzyme as methyltransferase for DNA (this work) and tRNA^28^ points to an adaptative evolutionary process in malaria parasites. Our findings expand the plasmodial epigenetic marks to three cytosine 5mC oxidative modifications and establish a direct link between variable oxygen environments encountered during the life cycle and the DNA cytosine modification pathway in malaria parasites.

## Materials and methods

### *P. falciparum* and mESC in vitro culture

*P. falciparum* 3D7-wildtype and Pf-DNMT2KO clones 1 and 2 parasites were maintained in culture and synchronized as previously described^29^. Parasites were routinely cultured at 5% normoxic O_2_ levels and for specific experiments at 20% hyperoxic O_2_ levels keeping CO_2_ levels well controlled at 5%. For genomic DNA preparation, parasites were first lysed using 0.15% saponin and parasite pellets were used to extract gDNA for digestion and hydrolysis.

mESC cells were cultured in standard mouse ES medium supplemented with LIF (1000 U/mL, Life Technologies) to prevent differentiation, and incubated at 37 °C, 5% CO2 as describe earlier^30^.

### Genomic DNA extraction

gDNA from parasites and mESC pellets was prepared using phenol-chloroform extraction. Briefly, 100 μL of DNA Extraction Buffer and 15μL of Proteinase K from Thermo Fisher (# EO0491) were added to 100ng of gDNA. The tubes were mixed and incubated at 55°C overnight. Equal volume of Phenol/Chloroform (pH 8.0) was added to the digested sample. The tubes were then spun for 5 min at 15.000 rpm separating the phases into the aqueous and the organic. For each sample, 650 μL 70%EtOH and 200 μL of 7.5 M ammonium acetate (AcNH4), were added to each microcentrifuge tube and vortexed. The tubes were then placed in a −20°C freezer overnight to precipitate DNA. Once extracted, gDNA was treated twice with a mixture of RNAse A/ RNAse T1 at 10 ug/μL for 1 h with phenol extraction and ethanol precipitation steps between the two treatments to ensure the total elimination of RNA.

### gDNA Enzymatic Digestion Protocols

#### Benzonase

100 ng of gDNA was dissolved in 100 μL of 10 mM Tris-HCl buffer pH 7.9 containing 10 mM MgCl_2_, 50 mM NaCl, 5 μM BHT from Sigma (B1378-100 G), and 3 mM deferoxamine from Sigma (# BP987). Next, 3 U of benzonase (in 20 mM Tris-HCl pH 8. 0.2mM MgCl_2_, and 20 mM NaCl), 4 mU phosphodiesterase I from Sigma (# P3243-1VL), 3 U DNAse I, 2 mU of phosphodiesterase II from Sigma (# P9041-10 UN) and 2 U of alkaline phosphatase from Biolabs (# M0290) were added and the mixture was incubated at 37°C. Finally, the mixture was transferred in a microspin filter from Agilent Technologies (# 5190-5275) and recovered after spinning at 20°C for 10 min at 14 000 rpm.

#### Nuclease P1

We used a modified protocol from (Wang et al; 2011)^31^. 100 ng of gDNA was dissolved in a 10 μL of buffer containing 0.3 M AcONa pH 5.6, 10 mM ZnCl_2_, 3m M deferoxamine, and 1 mM EHNA. Next, 4 U of Nuclease P1 from Sigma (# N8630-1VL) (in 30 mM AcONa pH 5.3, 5 mM ZnCl_2_ and 50 mM NaCl) and 5 mU phosphodiesterase II from Sigma (# P9041-10UN) were added and the mixture was incubated at 37°C. After 4 h, 4 U of alkaline phosphatase from Biolabs (# M0290) and 5 mU of phosphodiesterase I from Sigma (# P3243-1VL) were added and the mixture was incubated for additionally 2h. Finally, the mixture was transferred in a microspin filter from Agilent Technologies (# 5190-5275) and recovered after spinning at 20°C for 10 min at 14 000 rpm.

#### NEB Nucleoside Digestion kit

100 ng of gDNA was digested according to the Biolabs New England manufacture using the Nucleoside Digestion kit (# M0649), with little modification on time incubation, extended to overnight at 37°C. Finally, the mixture was transferred in a microspin filter from Agilent Technologies (# 5190-5275) and recovered after spinning at 20°C for 10 min at 14 000 rpm.

#### DNA Degradase Plus kit

100 ng of gDNA was digested according to the Zymo Research manufacture using DNA Degradase Plus™ kit (# E 2020), with little modification on time incubation, extended to overnight at 37°C. Finally, the mixture was transferred in a microspin filter (3 kDa) and recovered after spinning at 20°C for 10 min at 14 000 rpm.

#### Nuclease S1

We used modified protocol from (Globisch et al., 2010)^32^. 100 ng of gDNA was heated to 100°C for 5 min to denature the DNA and rapidly cooled on ice. Buffer A (10 μL, 300 mM ammonium acetate, 100 μM CaCl_2_, 1 mM ZnSO_4_, pH 5.7) and nuclease S1 from Sigma (# N5661-50 KU) (80 U, *Aspergillus oryzae*) were added to the mixture and incubated for 3 h at 37°C. Addition of buffer B (12 mL, 500 mM Tris-HCl, 1 mM EDTA), Antarctic phosphatase from NEB (# M0289S-10 units), snake venom phosphodiesterase I (0.2 units, *Crotalus adamanteus* venom) from Sigma (# P3243-1VL) and incubation for further 3 h at 37°C completed the digestion. Finally, the mixture was transferred in a microspin filter from Agilent Technologies (# 5190-5275) and recovered after spinning at 20°C for 10 min at 14 000 rpm.

### Quantification of DNA cytosine modifications by LC-MS/MS

Analysis of global levels of 5mdC, 5hmdC, 5fdC and 5cadC (herein abbreviated 5mC, 5hmC, 5fC and 5caC) was performed on a Q exactive mass spectrometer (Thermo Fisher Scientific). It was equipped with an electrospray ionisation source (H-ESI II Probe) coupled with an Ultimate 3000 RS HPLC (Thermo Fisher Scientific). DNA digestions were injected onto a Thermo Fisher Hypersil Gold aQ chromatography column (100 mm * 2.1 mm, 1,9 um particle size) heated at 30°C. The flow rate was set at 0.3 mL/min and run with an isocratic eluent of 1% acetonitrile in water with 0.1%formic acid for 10 minutes. Parent ions were fragmented in positive ion mode with 10% normalized collision energy in parallel–reaction monitoring (PRM) mode. MS2 resolution was 17,500 with an AGC target of 2e5, a maximum injection time of 50 ms and an isolation window of 1.0 m/z. The inclusion list contained the following masses: dC (228.1), 5mdC (242.1), 5hmdC (258.1), 5fdC (256.1) and 5cadC (272.1). Extracted ion chromatograms of base fragments (±5ppm) were used for detection and quantification (112.0506 Da for dC; 126.0662 Da for 5mdC; 142.0609 for 5hmdC; 140.0449 for 5fdC and 156.0399 for 5cadC). Calibration curves were previously generated using synthetic standards in the ranges of 0.2 to 50 pmoles injected for dC and 0.02 to 10 pmoles for 5mdC, 5hmdC, 5fdC and 5cadC. Results are expressed as a percentage of total dC

### Yeast culture and strain construction

Standard methods were used for cultivation and manipulation of yeast strains^33^. Epitope-tagging was performed as described previously^34^. In order to express Dnmt2^*pf*^-SNAP, the coding sequence of the SNAP-tag was PCR amplified using the source plasmid pAct –SNAP-3XGGGs-GW gift from Anne Plessis Lab and the primers GCGGGTACCATGGACAAAGACTGCGAAATG and GCGCTCGAGTCATTAGCCCAGCCCAGGCTTGCCCAG. The 2,713 kbp HindIII XhoI fragment containing the Dnmt2-SNAP fused sequence was excised and inserted into pRS416 plasmid gift from GF Richard lab. This vector is a centromeric (*i.e*., low-copy number) plasmid designed for the expression in yeast of hybrid proteins with SNAP fluorescent protein at the N-terminus.

### SNAP immunofluorescence assay

For the evaluation of yeast transformation, the commercially available fluorescent ligand SNAP-Cell TMR-Star (New England Biolabs) was used.

For image acquisition, an epifluorescence microscope (Leica Microsystems, Germany) equipped with an oil immersion lens (1.4 NA; 100 x; DM200 LED; Leica Microsystems), a N3 filter cube (excitation: 546/12 nm; emission: 600/40 nm) and a GFP filter cube (excitation: 450/90 nm; emission: LP 590 nm) was employed.

### Dot Blot Assay

20 μg of gDNA was denatured at 100°C for 10 min. Samples were rapidly chilled for 5 minutes on wet ice and then applied to a positively charged nylon membrane under vacuum using a 96-well Dot Blot Hybridization Manifold (Harvard Apparatus Limited, Holliston, MA, USA). The membrane was washed twice in 2 x SSC buffer and UV-crosslinked. The membrane was probed with anti-bodies specific to 5mC (Active motif # 39649; dilution factor 1:1. 000). To control loading, the membrane was probed with a rabbit poly-clonal antibody raised against single-stranded DNA (Demeditec Diagnostics, Kiel, Schleswig-Holstein, Germany). Following treatment with enhanced chemiluminescence substrate, membranes were scanned on an Image Quant LAS 4000 (GE Healthcare, Little Chalfont, Buckinghamshire, UK) imaging station.

### DNA content quantification by flow cytometry

Parasites cultured at 5% or 20% O_2_ were fixed in 4% PFA + 0,0075% glutaraldehyde in 1x PBS. Fixed cells were then stained with 2X SYBR-Green and the mean fluorescence intensity was measured using a BD Fortessa FACS instrument and analyzed using FlowJo software.

### Statistics

Statistical analysis was carried out a 2-way ANOVA significance test, unless otherwise specified.

## Results

### An optimized mass spectrometry-based protocol for the detection of DNA cytosine modifications in AT-rich gDNA

In order to accurately investigate the levels of DNA cytosine epigenetic marks in the highly AT-rich genomic DNA (gDNA) of *P. falciparum* using LC-MS/MS, we compared the gDNA digestion and hydrolysis protocol using five different methods: Nuclease P1, Benzonase, NEB Nucleoside Digestion kit, Degradase Plus kit and Nuclease S1 (Figure 1). All methods showed similar levels of 5mC and oxidative cytosine forms in gDNAs from mouse Embryonic Stem Cells (mESC) (Figure 1A). However, in the case of malaria parasites, the detected levels of 5hmC varied dramatically dependent on the method used to digest and hydrolyze gDNA. The levels of 5hmC in *Plasmodium* gDNA samples treated with Nuclease P1 are up to 8-fold higher compared to all other hydrolysis methods (Figure 1B and Figure S1). Thus, our data show that Nuclease P1 (NP1) digestion is by far the most efficient enzyme to detect 5hmC in an extremely AT-rich organism. Therefore, the NP1 hydrolysis method was adopted to explore cytosine methylation marks in *P. falciparum.*

**Figure 1:**
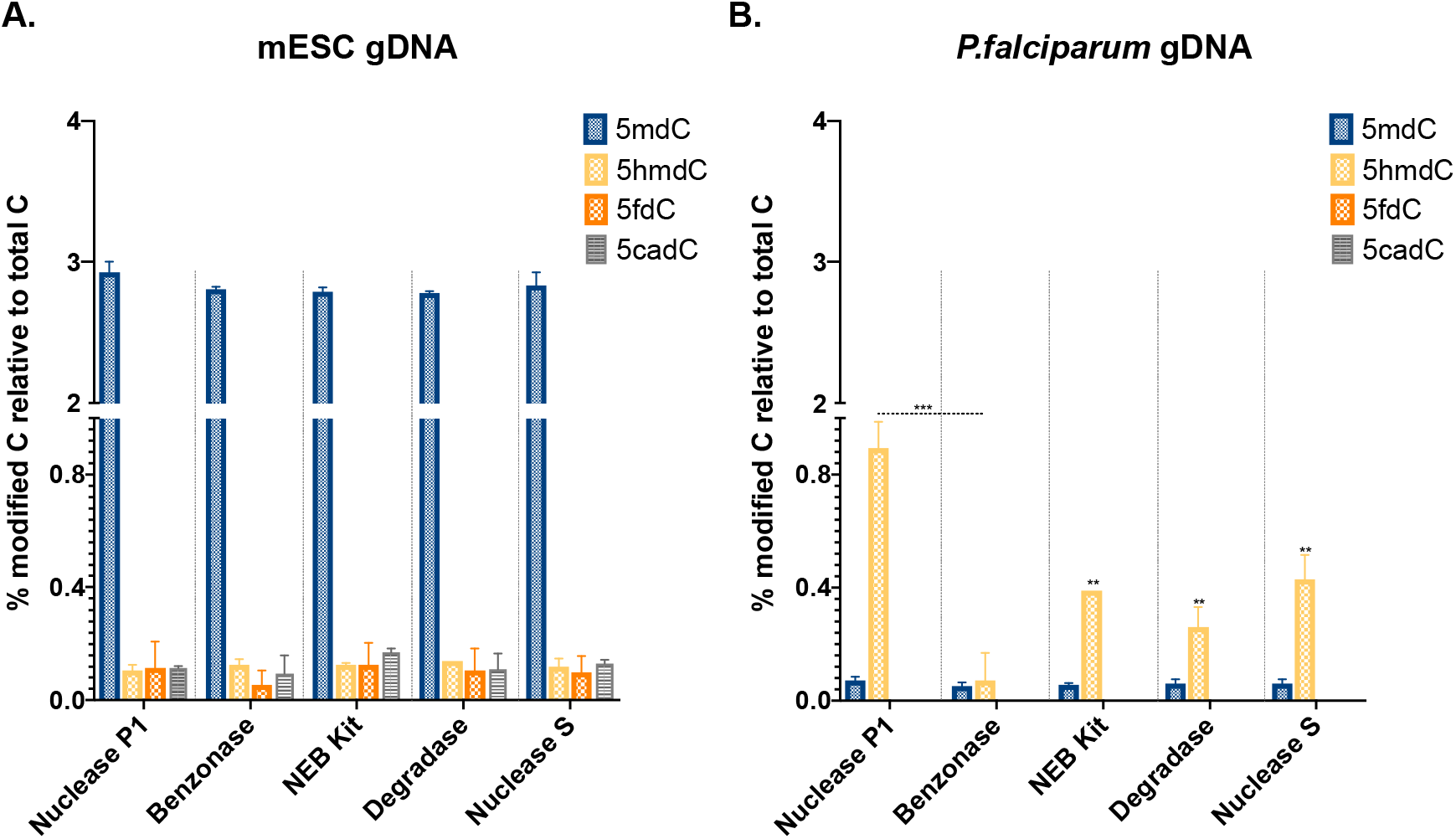
Comparative cytosine modification analysis by LC-MS/MS of gDNA from *P. falciparum* and mouse embryonic stem cells (mESC) using different digestion methods. Genomic DNA from mESC cells (A) and *P. falciparum* (B) was digested using five different protocols: Nuclease P1(NP1), Benzonase, NEB Nucleoside Digestion kit (NEB kit), Degradase kit (Degradase) and Nuclease S. Quantitative data analysis is shown as percentage of modified deoxycytidines relative to the total number of deoxycytidines in the tested sample and represents the mean (± SD) of two independent replicates for each condition (n=2). Statistical analysis was carried out using a 2way ANOVA test. P Value (Benzonase/NP1)=0,0007(***); P Value (NEB kit/NP1)=0,0048(**); P Value (Degradase/NP1)=0,002(**) and P Value (Nuclease S/NP1)=0,0065(**).

### DNA cytosine modifications are dynamically regulated during the 48-hour blood stage cycle of *P. falciparum*

Parasites were synchronized and harvested at the ring (10-15 hours post invasion; hpi), trophozoite (24-30hpi) and schizont stages (35-40hpi) to determine how the levels of 5mC and its oxidative derivatives vary across the intraerythrocytic developmental cycle (IDC) of *P. falciparum.* Using the LC-MS/MS protocol on DNA digested with Nuclease P1, we show that the levels of DNA cytosine modifications in malaria parasites dynamically change as the parasite transitions from the early ring stage to the mature schizonts in the infected red blood cells (iRBCs) (Figure 2A). While 5mC levels are very low in the ring (0.077% ± 0.015%) and schizont stages (0.13% ± 0.064%), we observe a drastic increase (10-fold) from the ring stage transition to trophozoites (0.727 % ± 0.265%) (Figure 2A). Surprisingly, 5hmC levels exhibit the opposite pattern. 5hmC is by far the predominant cytosine modification in rings (0.533% ± 0.352%) and schizonts (0.697%±0.072%) and decreases by 1.6-fold at the trophozoite stage (0.333%± 0.162%). In addition, the levels of 5fC and 5caC are extremely low in all three stages of the IDC with 5caC being only detectable in the schizont stages (0.03% ± 0.01%).

**Figure 2:**
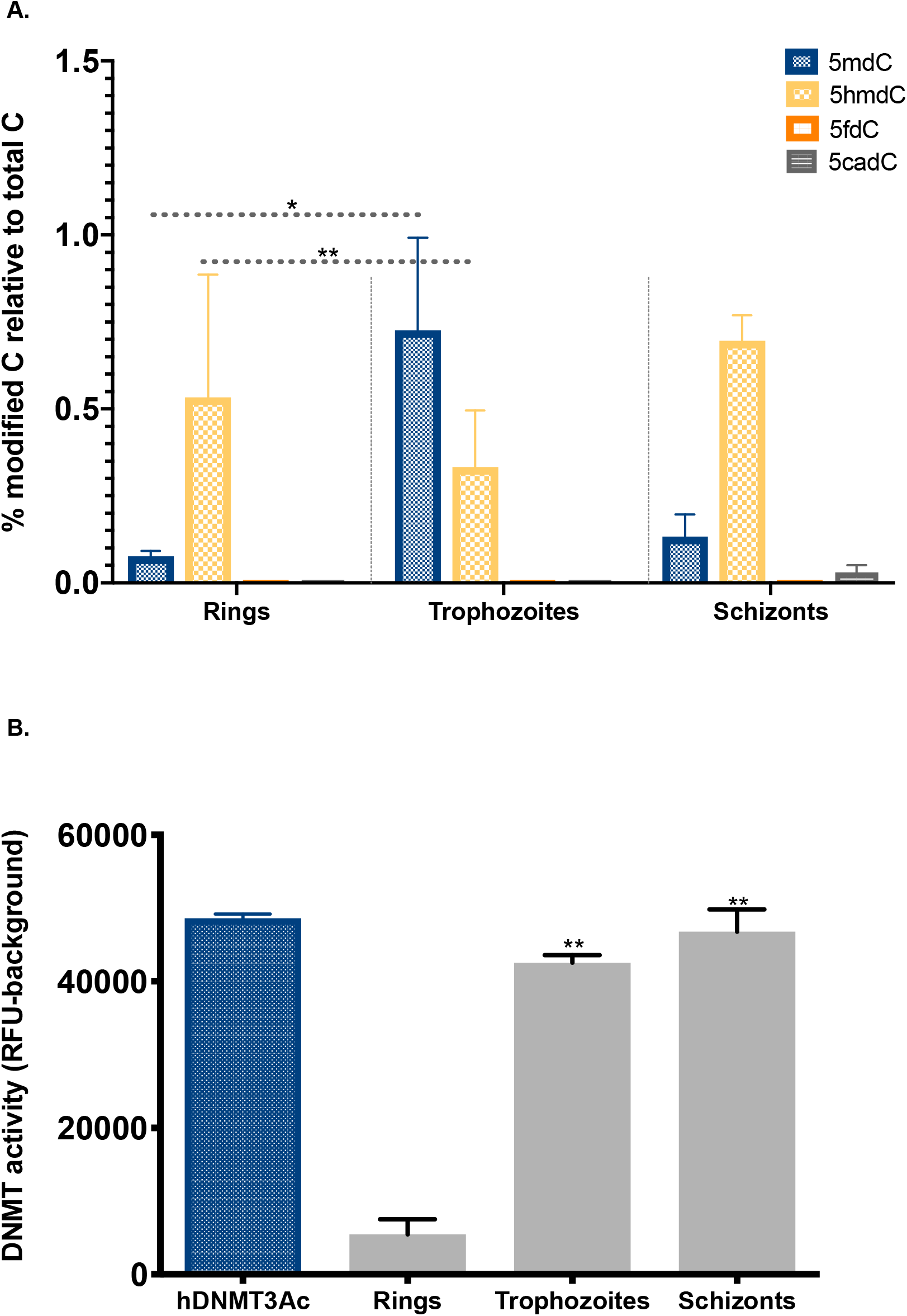
DNA cytosine methylation is dynamically regulated during the 48-hour *P. falciparum* asexual blood stage development. A. LC-MS/MS-based quantification of 5mdC, 5hmdC, 5fdC and 5cadC in gDNA of synchronized ring, trophozoite and schizont stages. Data is shown as percentage of modified nucleoside relative to the total number of deoxycytidines in each sample and represents the mean of three independent replicates for each stage (n=3). B. DNA methyltransferase (DNMT) activity assay using nuclear extracts from synchronous rings, trophozoite and schizonts. Data is shown as the RFU (sample)-RFU (background) and represents the mean (± SD) of two independent replicates for each stage (n=2). Human recombinant DNMT3aC was used as internal positive control. Statistical analysis was carried out using a 2-way ANOVA for the LC-MS/MS data and a t-test for the DNMT activity data. P Value <0.05 = *; P Value <0.005 =**.

Along with LC-MS/MS, we used a validated *in vitro* enzymatic assay^35^ in order to quantify DNA methylation activity (DNMT) in nuclear extracts prepared from the three stages of the IDC. Human recombinant DNMT3aC is used as internal positive control and the data is presented as relative fluorescence units (RFU) (sample)-RFU (background signal) (Figure 2B). Our results show that DNA methylase activity is very low in ring stages (5000 ± 200 RFU) with a sharp increase in activity (8-fold) in trophozoites (40000 ± 200 RFU). The high activity signal is maintained in schizonts (Figure 2B). The DNMT activity assay data corroborates our LC-MS/MS findings showing the sharp increase of DNA methylation in the trophozoite stages of *P. falciparum.* Altogether, our data shows that DNA cytosine methylation is very dynamic in malaria blood stages and suggests a tight developmental regulation of 5mC and the oxidative forms throughout *P. falciparum* IDC.

### Variable oxygen levels modulate the DNA cytosine methylation epigenetic landscape in *P. falciparum*

We next wanted to evaluate how different physiological oxygen environments might affect the levels of DNA cytosine modifications in the parasites asexual blood stages. To this end, we analyzed the hyperoxic (atmospheric) 20% O_2_ condition mimicking the levels of oxygen that the parasite encounters during *Anopheles* mosquito vector development. As control, the normoxic 5% O_2_ applied in standard *P. falciparum in vitro* cultures was used. Synchronous ring stage parasites were exposed to both 5% or 20% O_2_ levels and trophozoites, schizonts and rings were harvested for each condition during the next 72 hours. Giemsa-stained blood smears showed no morphological differences between trophozoites, schizonts and rings parasites cultured in normoxic or atmospheric O_2_ conditions (Figure S2). In addition, SYBR green-based DNA quantification showed that O_2_ levels do not impact parasite DNA content of blood stages (Figure S2) as suggested by previous studies^25^.

Importantly, LC-MS/MS-based DNA modifications quantification shows a shift of 5mC towards its oxidative forms in parasites cultured under hyperoxic 20% O_2_ levels (Figure 3). Notably, exposure of the parasites to hyperoxic O_2_ levels led to a significant increase in the abundance of 5caC in all three stages of the IDC (rings, trophozoites and schizonts) (Figure 3A, B and C) indicating an increased oxidation of 5hmC. The shift in the DNA methylation/oxidation balance is very clear at the trophozoite stage. 5mC levels are significantly decreased (14-fold) from (0.94% ± 0.11%) in parasites cultured under the control 5% O_2_ levels to (0.065% ± 0.017%) in those under 20% O_2_. The decrease of 5mC is accompanied by an increase in 5hmC (2.5-fold) from (0.3% ± 0.1 %) to (0.74% ± 0.04%), along with the significant increase in 5caC (16.5-fold) from (0.01%) to (0.17 ± 0.07%) (Figure 3B). Altogether, the LC-MS/MS data strongly support a direct influence of oxygen O_2_ concentration on the ratio of 5mC and its oxidative derivatives, which may allow the malaria parasite to adapt its epigenetic cytosine landscape to variable host environments.

**Figure 3:**
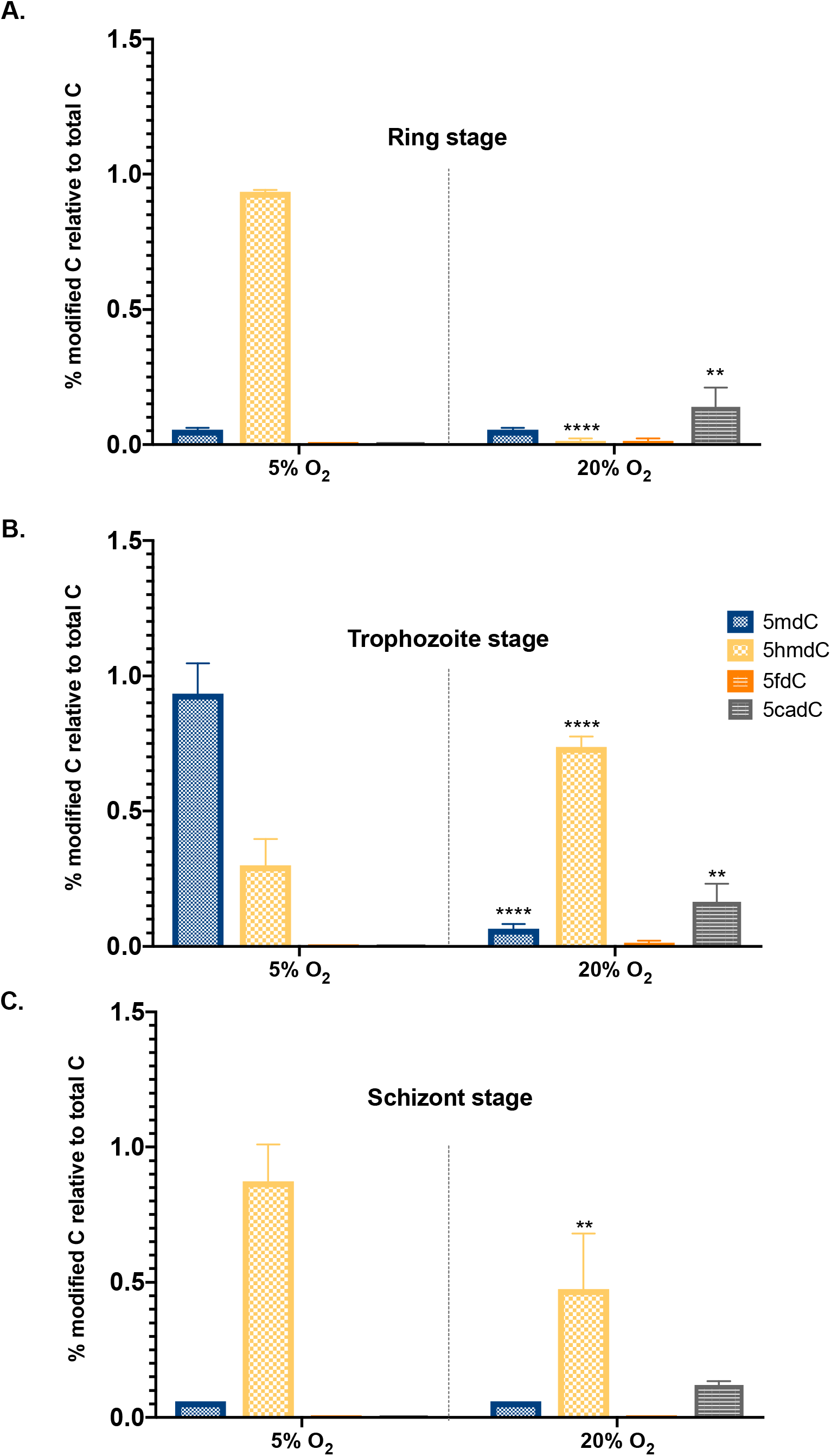
Effect of variable oxygen concentrations on gDNA cytosine modifications. DNA was isolated from *in vitro* cultured parasites cultured at different oxygen levels. Data is shown as percentage of modified deoxycytidines relative to the total number of deoxycytidines in each sample and represents the mean (± SD) of two independent replicates rings (A) and schizonts (C)(n=2) and four independent replicates for the trophozoite (B)(n=4). Statistical analysis was carried out using a 2way ANOVA test. P Value <0.0001 = ****; P Value <0.005 =**.

### DNA cytosine methylation in *P. falciparum* is determined by DNMT2

Given that *P. falciparum* belongs to a small group of DNMT2-only organisms^27,36^, we investigated if the loss of Pf-DNMT2 impacts the DNA cytosine methylation repertoire of the parasite. To this end, we used trophozoites genomic DNA from two Pf-DNMT2KO clones (cl1 and cl2) that we generated previously^28^. The LC-MS/MS analysis show that the levels of 5mC are drastically decreased to background levels in the Pf-DNMT2KO clones (0.03%) when compared to the 3D7 control strain (0.620% ± 0.269%) (Figure 4B). Importantly, 5hmC levels were also dramatically decreased in the two Pf-DNMT2KO clones (0.009%) when compared to their 3D7 counterparts (0.42% ± 0.085%). In addition, loss of 5mC and 5hmC was accompanied by a 20-fold decrease of 5caC in Pf-DNMT2KO-cl1 and Pf-DNMT2KO-cl2 (Figure 4B trophozoite and Figure S3 for rings and schizont stage).

**Figure 4:**
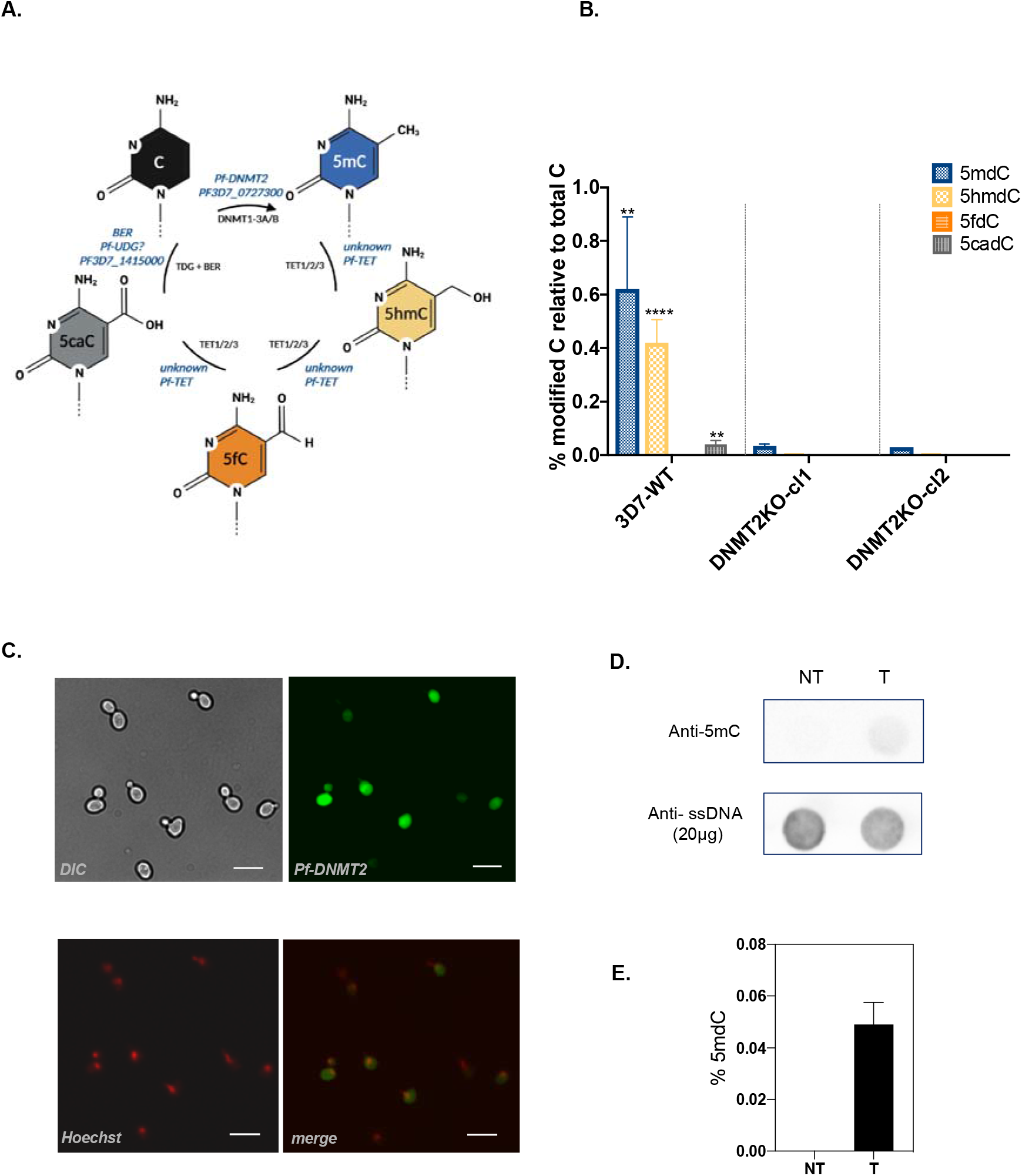
DNA cytosine methylation marks in *P. falciparum* are dependent on the activity of Pf-DNMT2. A. Schematic representation summarizing the DNA methylation/demethylation pathway as established in model organisms (black font) compared to what is known about the DNA methylation/demethylation machinery in *P. falciparum* (blue font). Pf-DNMT2: *P. falciparum* DNA methyltransferase 2; TET: ten eleven translocation enzymes; BER: base excision repair machinery; TGD/UDG: thymine/uracil-DNA glycosylase. B. LC-MS/MS-based quantification of 5mdC, 5hmdC, 5fdC and 5cadC in gDNA from 3D7-wild type (WT) parasites *versus* gDNA from the two Pf-DNMT2KO clones cl1 and cl2. Data is shown as percentage of modified deoxycytidines relative to the total number of deoxycytidines in each sample and represents the mean (± SD) of two independent replicates of each strain(n=2). C. DNMT2-SNAP expression in *S. Cerevisiae* cells. Scale bar=10 μm. DIC: differential interference contrast; Hoechst: nuclear staining. D. Immuno dot-blot analysis of 5mC in gDNA from yeast cells non-transformed (NT) or transformed (T) with Pf-DNMT2-SNAP plasmid. E. LC-MS/MS-based quantification of 5mdC in gDNA from yeast cells nontransformed (NT) or transformed (T) with Pf-DNMT2-SNAP plasmid. Statistical analysis was carried out using 2way ANOVA. P Value <0.0001 = ****; P Value <0.005 =**.

To confirm the function of Pf-DNMT2 as a DNA methyltransferase we expressed the full-length protein in the budding yeast *Saccharomyces cerevisiae (S. Cerevisiae*), which lacks any of the known DNMTs^37^ (Figure 4C). The 5mC content in transformed and non-transformed yeast cells was first assessed using a monoclonal antibody specific to 5mC. Methylated DNA is detectable only in Pf-DNMT2 transformed yeast cells (figure 4D). LC-MS/MS analysis of digested DNA isolated from non-transformed yeasts shows no detectable 5mC as expected. However, in Pf-DNMT2 transformed yeast cells the levels of 5mC increased up to 0.06% of total cytosines (Figure 4E). Taken together, our data clearly show that Pf-DNMT2 (PF3D7_0727300) is an active DNA methyltransferase responsible of cytosine methylation in the gDNA of *P. falciparum* (Figure 4A). Importantly, the LC-MS/MS data also shows that the presence of 5hmC, 5fC and 5caC in *P. falciparum* is completely dependent on 5mC as previously shown for other model systems^14,15,20,38^. In addition, our results infer the presence of a yet unknown TET-like enzyme in *P. falciparum* that triggers the oxidation of 5mC into 5hmC and further into 5fC and 5caC (Figure 4A).

## Discussion

*P. falciparum* is among the most AT-rich genomes known to date with an overall A+T composition of 80.6%, which rises to ~90% in introns and intergenic regions^39^. Given this unusual base composition, we compared different nuclease digestion procedures into single nucleosides prior LC-MS/MS analysis of cytosine modifications in genomic DNA. We observed striking differences in the digestion efficacy of 5hmdC between the distinct nucleases. With Nuclease P1 (NP1) 5hmdC levels are up to 8-fold higher compared to all others digestion methods, highlighting an important difference between AT-rich genomes and organisms with a balanced nucleotide composition. AT-rich DNA sequences display features that could account for the observed low efficacy of digestion of the oxidized DNA cytosine modifications. For example, AT-rich DNA stretches can form local curvature^40^ that may exert an influence on the interaction of nucleases with DNA. Furthermore, 5hmC was shown to enhance DNA flexibility^41^, a feature that could change the affinity of certain type of nucleases for 5hmdC containing DNA.

We thus used nuclease P1 for digestion of gDNA to investigate the plasmodial DNA cytosine modifications landscape during the blood stage development using LC-MS/MS. Our results show several features not observed previously in other organisms. The ratio between 5mC and the three oxidized cytosine forms is rather unusual when compared to higher eukaryotes (Figure 1A/B). Furthermore, we observe dynamic changes of 5mC levels during the 48-hours blood stage parasite cycle. For most of the time, we detect only very low levels of 5mC (0.07-0.1%) and high levels of 5hmC (0.5-0.9%). But during the short period of parasite proliferation (trophozoite stage), the 5mC/5hmC ratio is inverted (Figure 2A). The reason for the observed dynamic changes remains elusive. While ring stages circulate in the blood, trophozoite stages start to be sequestered in capillaries of host organs. It is tempting to speculate that the change in parasite tropism during blood stage development may be linked to increased DNA methyltransferase activity or reduced DNA demethylation activity. Further studies are necessary to explore the underlying molecular basis. The levels of 5fC and 5caC remain very low in all three stages (standard *in vitro* culture condition), a situation similar to what has been reported in other eukaryotes.

Previous work has reported relatively low 5mC levels in *P. falciparum* genomic DNA^23,24^. In addition, LC-MS/MS analysis did not detect 5hmC but pointed to a cytosine base modification with unknown biochemical identity^23^. The digestion protocol used in this report was based on the benzonase nuclease. It is important to note that the unusual base reported earlier was also detectable in our nuclease digestion protocols (Figure S1). We observed that this cytosine form was predominant in *P. falciparum* gDNA hydrolyzed with benzonase but present only at very low levels when the DNA was digested with Nuclease P1 (Figure S1). We conclude that the choice of nuclease for the digestion of AT-rich DNA is important to maximize the liberation of oxidative cytosine nucleosides.

Experimental evidence from the cancer field demonstrated that reduction of oxygen levels in tumors is associated with a reduced ability to produce 5hmC^22^. Since *P. falciparum* is exposed to fluctuating oxygen concentrations during the life cycle, we investigated if malaria parasite cytosine modification marks respond to physiological changes in oxygen levels. Our results demonstrate that host-dependent oxygen concentrations result in important alterations in the cytosine modification landscape of *P. falciparum.* We observed a shift towards oxidative derivatives of 5mC at higher oxygen levels (Figure 3). DNA cytosine modifications are now considered to function as distinct epigenetic marks with distinct profiles in tissues/organs in higher eukaryotes^38^. A recent study used anti 5hmC antibodies to immunoprecipitate DNA regions enriched for this cytosine mark (hmeDIP-seq) of gDNA of *P. falciparum.* The presence of hydroxymethylated cytosine in gene bodies positively correlated with transcript levels in blood stages^23^. We hypothesize that the observed dynamic changes of the cytosine modifications may contribute to the malaria pathogen adaptation to changing environments, in particular during *P. falciparum* development within the mosquito vector.

TET enzymes, which catalyze the oxidative cytosine forms depend on oxygen as co-substrate^15,21^, may sense changes in oxygen levels and change the basal cytosine modification level toward oxidative cytosine forms. For example, TET1 activity as well as 5hmC levels are decreased in low oxygen conditions leading to the inhibition of the differentiation of mouse embryonic stem cells (mESC)^42^. Although TET-like activity was previously detected in nuclear extracts of *P. falciparum* using a TET hydroxylase activity kit (Abcam) no obvious ortholog of mammalian TET enzymes could be identified to date^23^. Further studies are needed to identify TET-like enzymes in malaria parasites and to explore the role of the oxidized cytosines as potential epigenetic marks.

An obvious question is if the identified oxidative cytosine forms (5hmC, 5fC, 5caC) depend on 5mC as a precursor or are formed by oxidative lesions. *P. falciparum* is part of a small number of organisms called DNMT2-only, given the absence of genes coding for the canonical DNA methyltransferase genes DNMT1 and DNMT3^36^. A recent study reported a *P. falciparum* Pf-DNMT2 KO parasite line, and this work confirmed the well-established function of DNMT2 as tRNA aspartic acid (GTC) cytosine methyltransferase^28^. These KO parasites show no growth defect but are significantly more sensitive to various stressor and respond to specific stress situations by 8-fold higher transmission stages or higher drug sensitivity, respectively. This study also observed a reduction of 5mC in gDNA of KO parasites, however, given the very low level reported for 5mC (0.06%) in the investigated schizont stage, it remained unclear if the observed reduction in 5mC can be attributed to the loss of Pf-DNMT2 or variations in the experimental procedure^28^. DNMT2-dependent DNA methylation is still disputed with contradictory results from distinct DNMT2-only organisms^27^. Here we demonstrate that DNMT2 has a dual function using tRNA and DNA as substrate in malaria parasites. Quantitative LC-MS/MS analysis demonstrate that the levels of 5mC are drastically decreased to background levels in the Pf-DNMT2KO clones (0.03%) when compared to the 3D7 control strain (0.62%) (Figure 4B). Furthermore, the identified oxidative cytosine forms are also reduced to background levels, inferring that 5mC is the precursor for 5hmC, 5fC and 5caC cytosine modifications.

In conclusion, this work demonstrates that plasmodial Pf-DNMT2 has a dual function in methylating tRNA and DNA. The 5mC methylation of gDNA is developmentally regulated during the blood stage development and the 5mC distribution *versus* oxidized forms can vary dramatically depending on environmental oxygen concentrations. Expanding the number of potential epigenetic DNA marks in this pathogen may open new avenues to understand the regulatory circuits that act during the life cycle stage.

## Acknowledgements

This work was supported by the Laboratoire d’Excellence (LabEx) ParaFrap [ANR-11-LABX-0024], the Agence Nationale pour la Recherche [Project EpiKillMal, ANR-20-CE18-0006]. This work was supported by DIM1Health 2019 equipment grant “EpiK” from Région Ile de France.

## Author contributions

Conceptualization A.S, E.H, S.M and B.A; Methodology: E.H, S.M and F.B; Investigation E.H and S.M; Analysis: E.H, S.M and F.B; Writing: E.H, S.M and A.S; Funding acquisition: A.S and P.B.

## Data availability

LC-MS/MS raw data is shown in supplementary tables S1 and S2.

## Supplementary figures legends

**Figure S1: LC-MS/MS chromatogram of** deoxycytidine (dC), 5-methyl-2’-deoxycytidine (5mdC), 5-hydroxymethyl-2’-deoxycytidine (5hmdC), 5-carboxy-2’-deoxycytidine (5cadC), and 5’-formyl-2’-deoxycytidine (5fdC) in A) mESC gDNA and B) *P. falciparum* gDNA according to the digestion protocol. Blue arrows show the expected 5hmdC peak while the red arrows indicate the shifted 5hmdC peak observed earlier^23^.

**Figure S2: *In vitro* cultured parasite growth under normal (5% O2) and high oxygen concentrations (20% O_2_).**

Giemsa staining (A) and DNA quantification (B) during the 48h blood stage cycle do not reveal any developmental changes between parasites cultured at 5%*vs* 20% O_2_.

**Figure S3: LC-MS/MS quantification of 5mdC, 5hmdC, 5fdC and 5cadC in gDNA of DNMT2KO parasite from ring (A) and schizont stage (B).**

Nucleoside quantification of 5mdC, 5hmdC, 5fdC and 5cadC in gDNA from 3D7-wild type (WT) parasites *versus* gDNA from the two Pf-DNMT2KO clones cl1 and cl2. Data is shown as percentage of modified deoxycytidines relative to the total number of deoxycytidine in each sample and represents the mean (± SD) of two independent replicates of each strain.

