## Supplementary figures for "Variable oxygen environments and DNMT2 determine the DNA cytosine epigenetic landscape of *Plasmodium falciparum*"

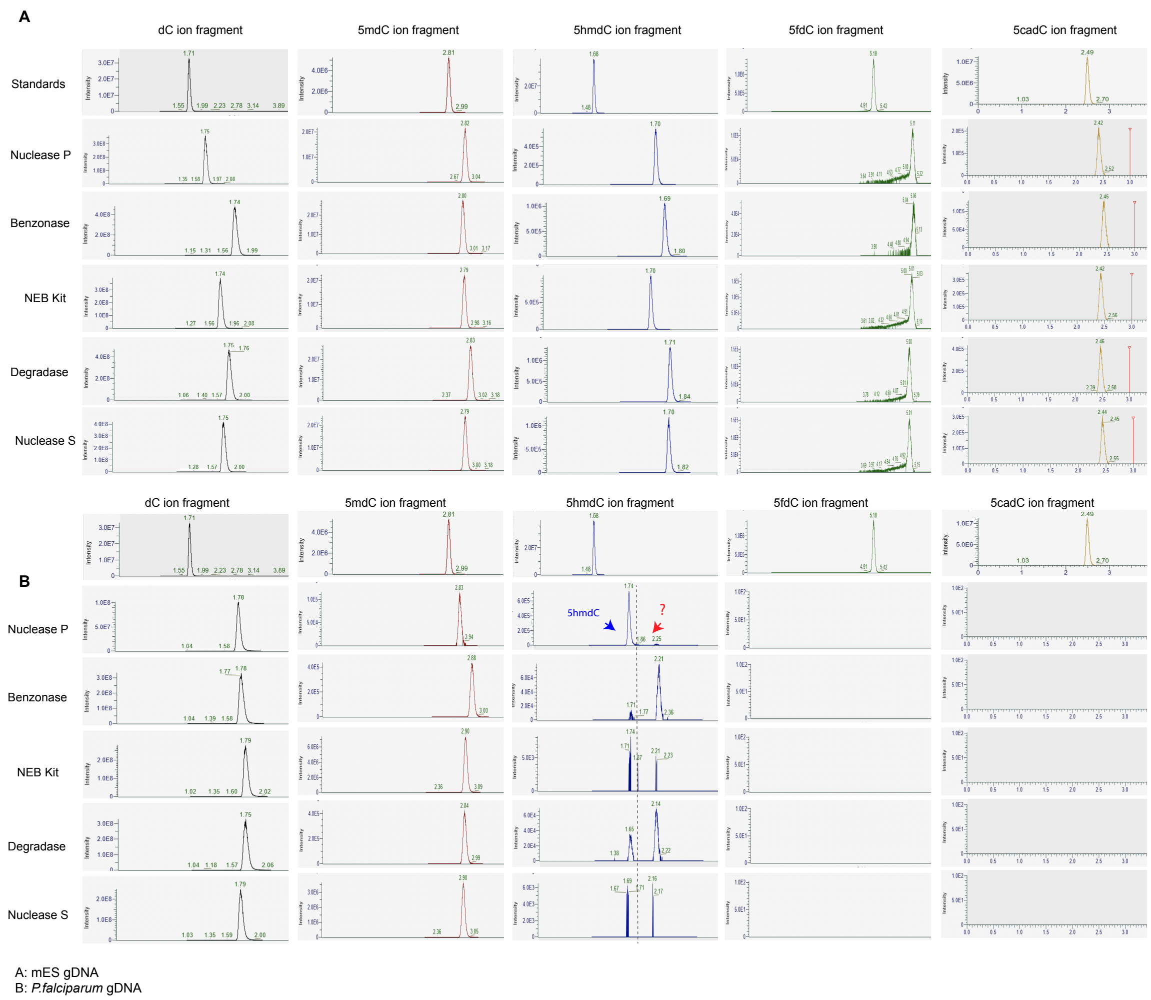


**Figure S1: LC-MS/MS chromatogram of** deoxycytidine (dC), 5-methyl-2’-deoxycytidine (5mdC), 5-hydroxymethyl-2'-deoxycytidine (5hmdC), 5-carboxy-2'-deoxycytidine (5cadC), and 5’-formyl-2'-deoxycytidine (5fdC) in A) mESC gDNA and B) *P. falciparum* gDNA according to the digestion protocol. Blue arrows show the expected 5hmdC peak while the red arrows indicate the shifted 5hmdC peak observed earlier[23].

A.

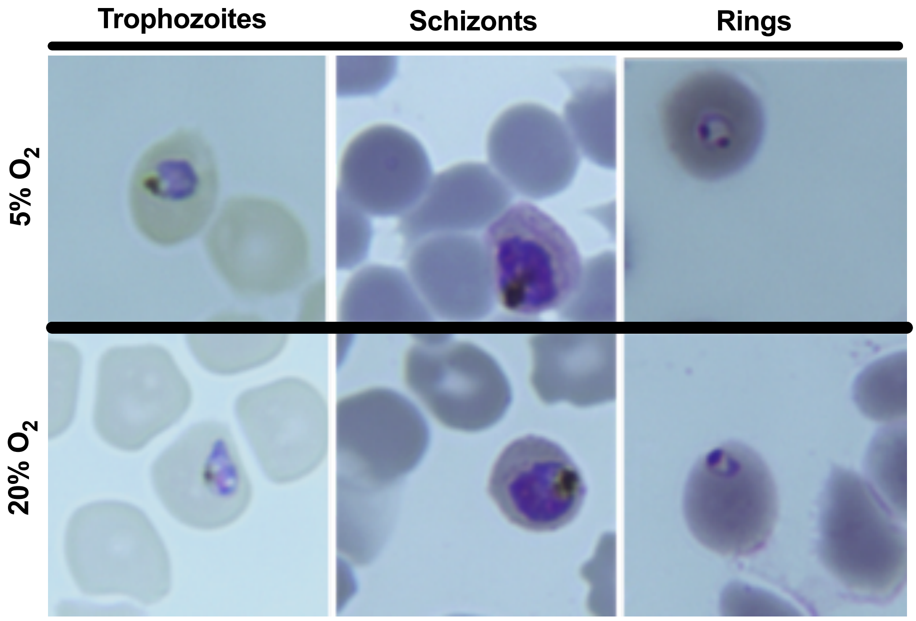
Trophozoites
Schizonts
Rings
5% O2
20% O2

B.

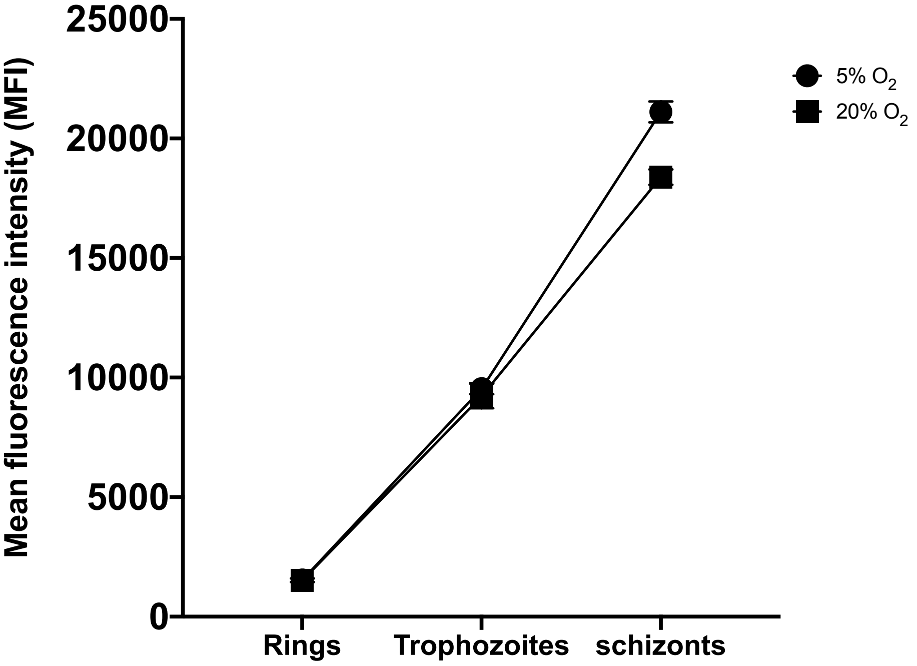
Mean fluorescence intensity (MFI)
25000
20000
15000
10000
5000
0
Rings
Trophozoites
schizonts
5% O2
20% O2

**Figure S2: *In vitro* cultured parasite growth under normal (5% $O_2$) and high oxygen concentrations (20% $O_2$).**

Giemsa staining (A) and DNA quantification (B) during the 48h blood stage cycle do not reveal any developmental changes between parasites cultured at 5% *vs* 20% $O_2$.

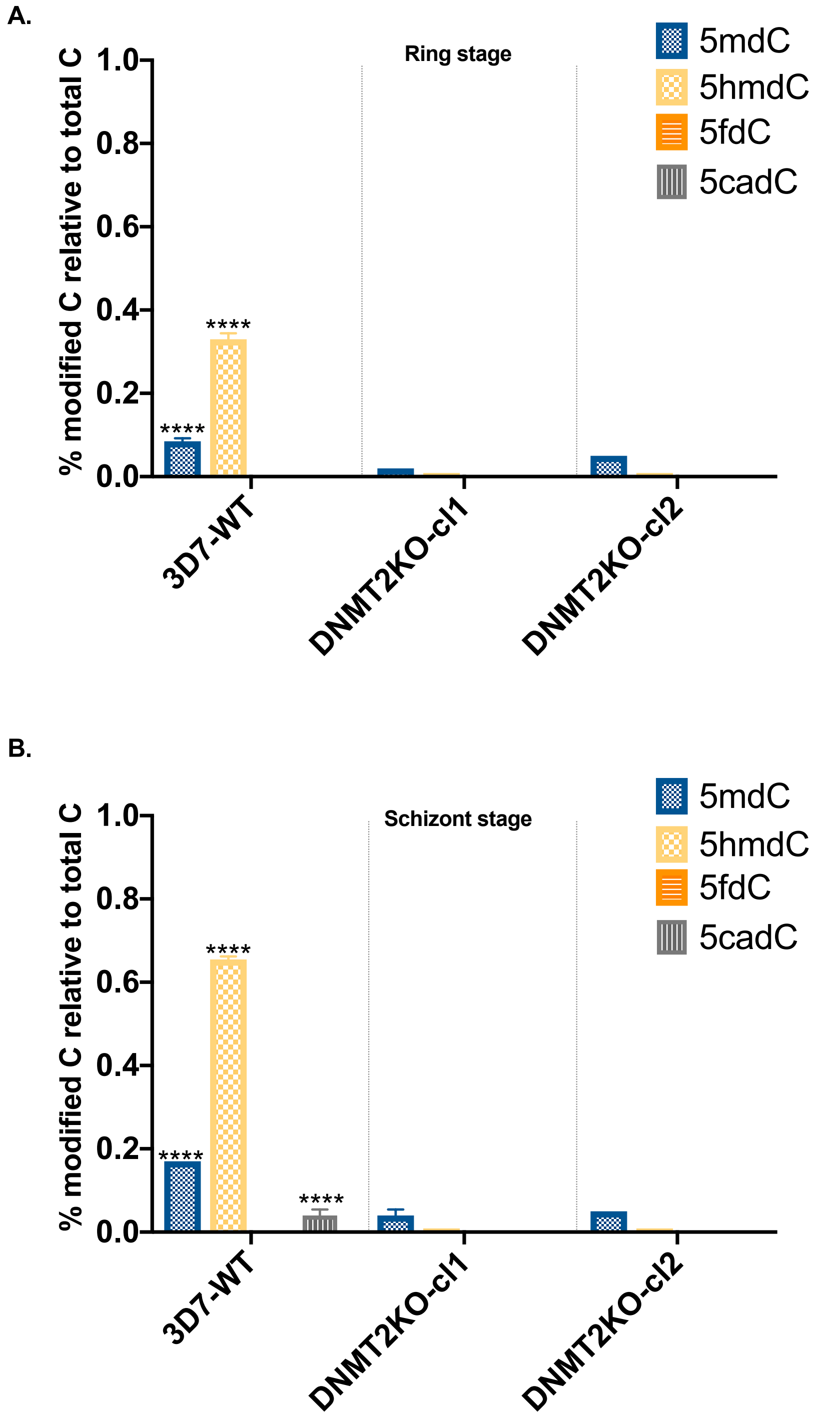

**Figure S3: LC-MS/MS quantification of 5mdC, 5hmdC, 5fdC and 5cadC in gDNA of DNMT2KO parasite from ring (A) and schizont stage (B).**

Nucleoside quantification of 5mdC, 5hmdC, 5fdC and 5cadC in gDNA from 3D7-wild type (WT) parasites *versus* gDNA from the two Pf-DNMT2KO clones cl1 and cl2. Data is shown as percentage of modified deoxycytidines relative to the total number of deoxycytidine in each sample and represents the mean (± SD) of two independent replicates of each strain.
